# Molecular determinants of large cargo transport into the nucleus

**DOI:** 10.1101/695080

**Authors:** Giulia Paci, Edward A Lemke

## Abstract

Transport of molecules between the nucleus and the cytoplasm is tightly regulated by the nuclear pore complex (NPC). Even very large cargoes such as many pathogens, mRNAs and pre-ribosomal subunits can pass the NPC intact. Compared to small import complexes, for such large cargoes >15 nm there is very little quantitative understanding of the mechanism for efficient transport, the role of multivalent binding to nuclear transport receptors via nuclear localisation sequences (NLSs) and effects of size differences. Here, we assayed nuclear import kinetics in cells for a total of 30 large cargo models based on four capsid-like particles in the size range of 17-36 nm, with tuneable numbers of up to 240 NLSs. We show that the requirements for transport scale non-linearly with size and obey a minimal cut off of functional import requiring more than 10 NLS in the lowest case. Together, our results reveal the key molecular determinants on large cargo import kinetics in cells.

## Introduction

Cargo transport across the nuclear envelope is a hallmark of eukaryotic cells and central to cellular viability. In a typical human embryonic kidney cell (HEK), more than 2000 nuclear pore complexes (NPCs) span the nuclear envelope. With ≈ 120 MDa in metazoans (Reichelt et al., 1990) and roughly half that weight in yeast, the NPC is probably the largest proteinaceous structure that can be found inside the cell. NPCs are the gatekeepers of nucleocytoplasmic transport and restrict access of cargoes with increasing size if bigger than the smooth cut off limit around 4 nm (Timney et al., 2016). However, if cargoes can bind to nuclear transport receptors (NTRs), as for example via a nuclear localization sequence (NLS), then the same cargo can rapidly enter into the nucleus. Kinetics of nuclear import mainly focusing on cargoes with 1 to 4 NLSs have been shown in several studies to follow a monoexponential behaviour (Kopito and Elbaum, 2007; Ribbeck and Gorlich, 2001; Timney et al., 2006).

NPCs are remarkable in the diversity of sizes of cargoes they can transport, ranging from viral import to nuclear export of pre-ribosomal subunits and mRNA complexes. Nuclear transport, especially for very large cargoes (> 15 nm), is also still an enigma for structural reasons. The NPC is formed by multiple copies of about 30 proteins, two thirds of which are folded proteins that assemble the NPC scaffold. The recent improvements in electron tomography (ET) paired with X-ray crystallography, has greatly expanded our knowledge on the organization of these folded components of the NPC (Kosinski et al., 2016; Lin et al., 2016; Szymborska et al., 2013; von Appen et al., 2015). This pore like scaffold is filled with a matrix of 10 intrinsically disordered proteins, known as FG nucleoporins (FG Nups) for their high enrichment in phenylalanine and glycine (F and G) residues. FG Nups have been estimated to be at concentrations in the mM range inside the NPC (Aramburu and Lemke, 2017; Frey and Gorlich, 2007). Our structural knowledge about the actual transport conduit compared to the scaffold is much lower, as its dynamic nature leads to a loss of electron density in the averaging process inherent to ET, leaving a ≈20 nm hole inside the structural map of the NPC tomogram. As transport of many large cargoes is believed not to damage the NPC, substantial amounts of FG Nups mass must be displaced in order to facilitate such transport events.

Despite its high biological relevance, nuclear transport of large cargoes is still poorly understood. In order to address this gap, we designed a set of large model cargoes in which we were able to control the cargo size and the number of exposed NLSs. We used a combination of spectroscopy and semi-automated microscopy assays to investigate the kinetics of nuclear import for cargoes ranging from 17 to 36 nm in diameter in permeabilised cells. Our results uncover the quantitative interdependence of cargo size and NLS number in a previously unexplored range, revealing an unexpected high NLS cut off informing on the molecular determinants of large cargo transport.

## Results and discussion

### A large cargo toolkit for nuclear transport studies

We first aimed to develop a set of model import cargoes with known size and tuneable number of NLSs (#NLSs) on their surface. Naturally occurring cargoes with multiple NLSs, such as proteasomes, pre-ribosomes, mRNA or RNA-protein complexes do not offer the possibility to control both properties reliably at the same time. Vice versa, for artificial large substrates, like quantum dots or gold nanoparticles, it can be challenging to tune size and #NLSs and extensive functionalization is typically required. Thus we turned to viral capsids, which are known to self-assemble from one or few proteins into large structures of fixed size. We screened the literature for capsid-like particles obeying the following criteria: i. Large-scale high yielding recombinant expression is possible in a host like E. coli. ii. Surface modification via a unique residue is possible. Hence we focused on systems where crystal and/or EM structures have been obtained and identified if single functional surface exposed cysteines existed or could be mutated with no impact on capsid assembly. iii. Stability of the capsid at physiological conditions. iv. To focus on rather uncharted territory, the capsid size was chosen to be larger than 15 nm and up to 36 nm, as this has been reported to be largest size of cargoes transported by the NPC. The following five icosahedral shaped capsids of different size were selected for this study (**Figure 1** and Supplementary **Figure S1**).

**Figure 1:**
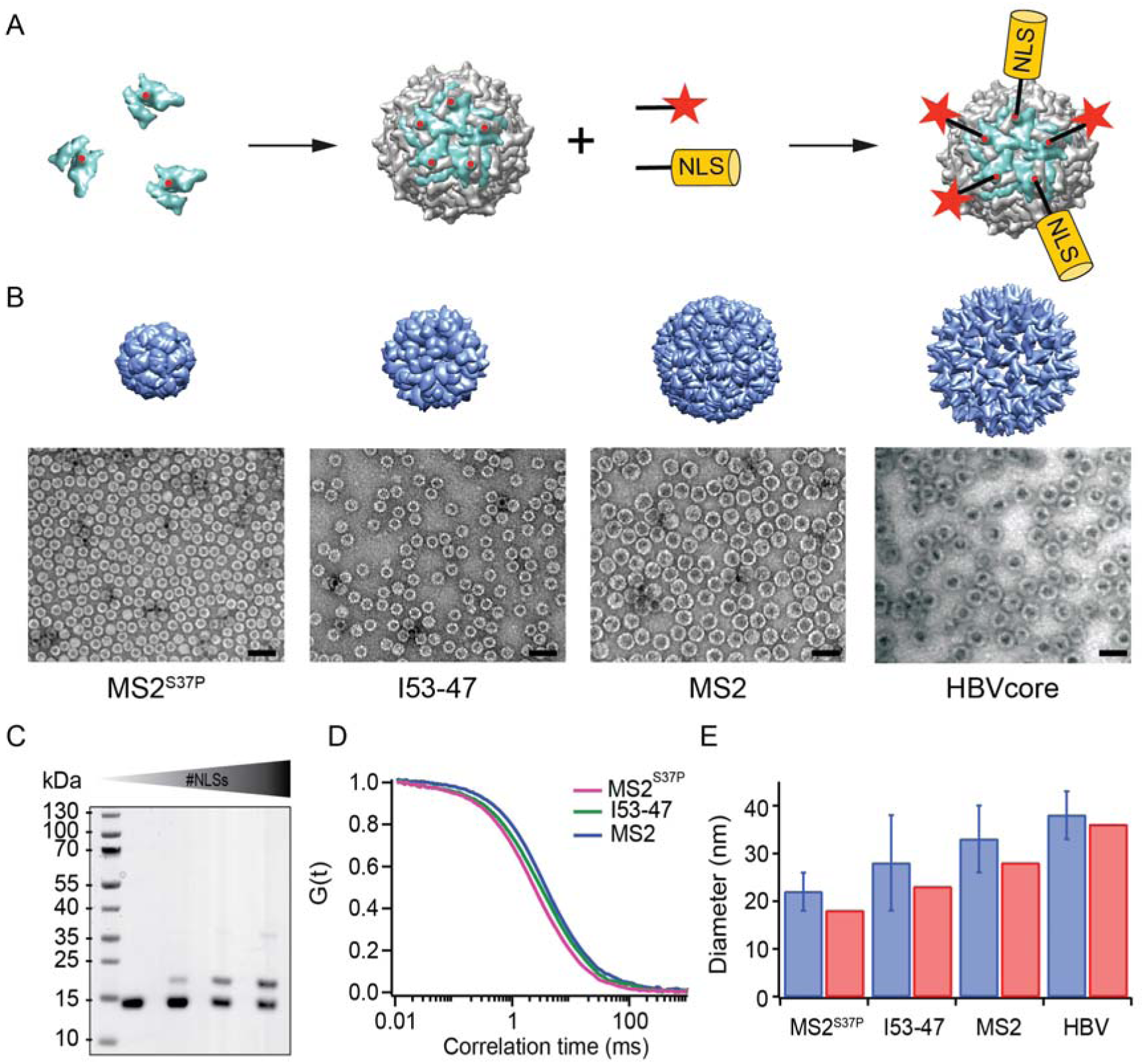
A large cargo “toolkit” for nuclear import studies. (A) Schematic representation of the mixed labelling reaction with maleimide reactive NLS peptide and maleimide reactive dye. The capsid protein, containing a cysteine mutation (in red), self-assembles into a capsid. The purified capsids are then labelled with a mixture of dye and NLS peptide, in different ratio according to the desired reaction outcome. (B) Capsid structures rendered in Chimera (Pettersen et al., 2004) (top) and EM images of the purified capsids (bottom). The scale bar corresponds to 50 nm. (C) SDS-PAGE gel of MS2^S37P^ samples with increasing number of NLS peptides attached (top band). The lower band corresponds to unlabelled capsid protein and capsid protein tagged with the dye. (D) Representative FCS autocorrelation curves for the MS2^S37P^, I53-47 and MS2 capsids. The curves were fitted with a diffusion model to calculate the capsid brightness and concentration. (E) DLS quantification of capsid diameters (blue bars) compared with reference values from literature and structural information (red bars).

MS2^S37P^ (diameter 17 nm): This capsid is derived from the bacteriophage MS2, formed by a single coat protein with a point mutation S37P. The coat protein assembles into dimers and then into 12 pentamers yielding an icosahedron with a total of 60 copies (Asensio et al., 2016). A cysteine mutation (T15C), that had previously been shown not to interfere with capsid assembly, was introduced to allow surface tagging via maleimide labelling (Peabody, 2003).

I53-47 (diameter 23 nm): This artificial capsid is derived from *de novo* designed capsids developed by the Baker’s lab (Bale et al., 2016). The I53-47 variant is formed by two different proteins (chain A and chain B), occurring in 60 copies each and organized in 12 pentamers and 20 trimers. A cysteine mutation exposed on the capsid surface was introduced in chain B (D43C), following recent work where different surface mutations were introduced in a similar capsid variant (Butterfield et al., 2017).

MS2 (diameter 27 nm): This capsid is derived from the wild type bacteriophage MS2 coat protein, which assembles into dimers and then into an icosahedron with 12 pentameric and 20 hexameric faces, to a total of 180 copies. Also here the same cysteine mutation as MS2^S37P^ (T15C) enabled tagging via maleimide labelling (Peabody, 2003).

I53-50 (diameter 27 nm): this capsid variant has a structure similar to the I53-47 described above, with a larger chain A protein that result in a capsid of increased size (Bale et al., 2016). The same cysteine mutation was introduced as for the I53-47.

Hepatitis B capsid (diameter 36 or 32 nm depending on isoform): this capsid is based on an assembly-competent truncated version of the HBV core protein (1-149). This truncation leads to higher levels of bacterial expression and to a predominance of the T=4 capsid with no obvious change in capsid morphology (Zlotnick et al., 1996). The core protein thus assembles into 36 nm capsids from 12 pentameric and 30 hexameric units, for a total of 240 copies. The truncation also results in the removal of the C-terminal native capsid NLS, enabling a complete control over number of exposed NLSs via surface engineering. A cysteine mutation (S81C) was introduced into an exposed loop of the core protein (c/e1 epitope) to allow surface tagging via maleimide labelling. Hepatitis B capsid, which can have 240 NLSs is frequently quoted as the largest cargo known to pass the NPC intact (Panté and Kann, 2002), and thus constitutes the upper limit of the cargoes we investigated.

After successful purification, the next step was to engineer the capsid surface with a fluorescent dye and with NLSs. As detailed in the methods, the use of tangential flow for sample concentration and buffer exchange turned out to be of highest practical relevance to purify preparative amounts of intact capsids for further labelling reactions (see Methods). We chose maleimide reactive dyes and a synthetic maleimide reactive NLS, with a sequence known to bind tightly to Importinα which binds to Importinβ via its IBB domain (Hodel et al., 2001). Capsids were labelled with suitable mixtures of dye and NLS peptide.

For the I53-50 capsid we observed non-specific labelling to both protein chains (see supplementary Figure S1), tentatively on a cysteine which according to our structural analysis should not have been surface exposed. Due to this labelling ambiguity, this capsid was excluded from further analysis.

**Figure 1** summarizes the labelling scheme used for all capsids and its charaterization. **Figure 1B** shows negative staining EM of capsids after purification and labelling, visualizing intact capsids with the expected diameter. For each capsid, it was crucial to determine its fluorescence brightness (i.e. how many dyes are attached to one capsid) as well as the number of NLSs. Dye/capsid number was determined via fluorescence correlation spectroscopy (FCS), a widely employed biophysical tool to probe brightness and concentration of a freely diffusing species (**Figure 1D**). FCS can also be used to estimate the size and size distributions (such as substantial contaminations of other species than intact capsids) of the capsids, which was found to be in line with the high purity indicated by the EM micrographs. Additional DLS (dynamic light scattering) studies were employed to further validate capsid diameter in solution and presence of intact capsids as the dominant species (**Figure 1E**). The number of NLSs was determined from gel shift assays, as labelled capsid monomers migrate substantially different then their unlabelled counterparts (**Figure 1C**). We note that the labelling with a synthetic NLS to which a dye was directly engineered was found to be impractical in preliminary experiments, as even minor contaminations of unreacted peptide resulted in ambiguous results in downstream analysis.

### Import kinetics of large cargoes are tuned by size and NLS numbers

The different labelled capsids (total 30) were subjected to nuclear import assays using the widely employed permeabilised cell assay (Adam et al., 1990). **Figure 2A** outlines the details of the experiments. In brief, mild digitonin treatment was used to permeabilise the plasma membrane of HeLa cells, leaving the nuclear envelope intact. In these conditions, functional nucleocytoplasmic transport can be reconstituted for a few hours by adding the key components of the transport machinery: Importinβ, Importinα, RanGDP, NTF2 (a NTR which allows recycling of RanGDP) and GTP to the cells. Intactness of the nuclear envelope as well as functional nuclear transport were always validated by a set of control experiments using fluorescently labelled dextran and model cargoes (see Methods). As shown in **Figure 2B** exemplarily for the MS2^S37P^, capsids labelled with NLSs showed an increased nuclear accumulation over time, indicative of functional nuclear import of capsids.

**Figure 2:**
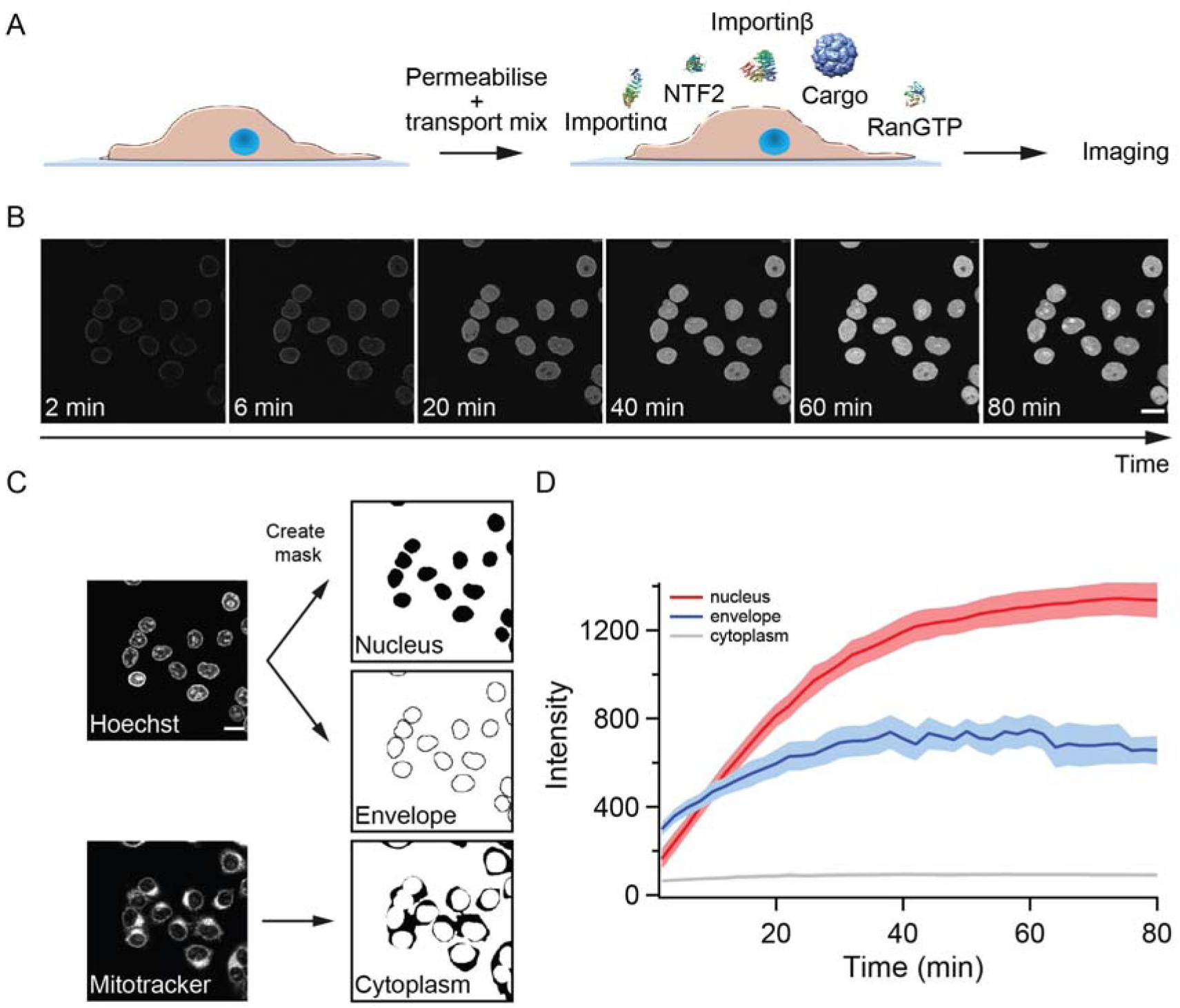
Pipeline for import kinetic experiments. (A) Scheme of the transport assay experiment: HeLa cells were permeabilised with digitonin and incubated with a transport mix containing the cargo of interest, nuclear transport receptors and energy. Confocal images were acquired in 12 different areas every 2 minutes, for 80 minutes in total. (B) Representative time lapse snapshots of cargo import (MS2^S37P^ capsid). The scale bar corresponds to 20 μm. (C) Overview of the image analysis pipeline for import kinetics experiments. Two reference stain images (Hoechst and MitoTracker) were segmented and used to generate three masks corresponding to the regions of interest: nucleus, nuclear envelope and cytoplasm. The masks were then applied to the cargo images to calculate the average intensity in the different regions. (D) Representative raw import kinetics traces for the three cellular compartments of interest. Curves depict the average fluorescence measure in the different regions; the shaded areas represent the standard deviation over 12 areas.

Experiments were performed on a semi-automated confocal microscope, recording time-lapse images over several cells and different field of views (shown error bars correspond to standard deviations between different FOV). Besides the nuclear signal, also nuclear envelope and cytoplasmic signals were recorded using suitable imaging masks (**Figure 2C** and methods for details). We took precautions to distinguish nuclear fluorescence from nuclear envelope fluorescence by eroding the nuclear mask to a region furthest away from the rim. This turned out to be important, as some capsids showed nuclear envelope targeting but no substantial accumulation into the nucleoplasm. In addition, this method enabled us to discriminate nuclear signal from sticking of capsids to the cytoplasm, which was observed in some cases.

**Figure 2D** summarizes the three kinetic traces that can be obtained from a typical experiment. In the representative experiment shown for a MS2^S37P^ capsid sample, the cytoplasmic fluorescence stayed constant, while nuclear envelope signal increased pointing to recruitment and accumulation of capsid at NPCs. The red curve shows the import kinetics of capsid into the nucleus. As discussed below, these could be fitted by a monoexponential kinetic model used to retrieve the saturation amplitude I_MAX_ and the half time to import 50% of the total capsids (T_1/2_). Supplementary **Figure S2** shows additional control experiments to establish that nuclear import rates are dependent on the substrate size and #NLSs and not on any of the component in the transport mix becoming limiting during the course of the experiment.

**Figure 3** shows representative nuclear import data for the three kinetically investigated capsids MS2^S37P^, I53-47 and MS2 (see Supplementary **Figure S3** for all dataset). The results for HBV capsids will be discussed in the next paragraph. Panel A displays typical confocal images for cargoes with different #NLSs and panel B shows representative nuclear kinetic traces extracted from semi-automated microscopy. In the absence of NLSs, all capsids localized to the cytoplasm and no targeting to the nuclear envelope nor accumulation into the nucleus was observed, in line with an Importin-dependent import pathway. With increasing #NLSs mounted onto the capsid surface, we observed progressive nuclear envelope targeting, and eventually efficient accumulation of cargo into the nucleoplasm. Strikingly, the #NLSs required to observe the same behaviour with different capsids scaled dramatically with cargo size, as can be seen by comparing e.g. the I53-47 sample image with 35 NLSs and the MS2 one with 86 NLSs. The observation of robust bulk import for all capsid constructs with sufficiently high #NLSs highlights another benefit of using viral capsids as large cargoes, as in a previous study using coated quantum dots (18 nm) only conditions were identified where rare events could be captured by advanced single molecule technologies but no bulk import (Lowe et al., 2010).

**Figure 3:**
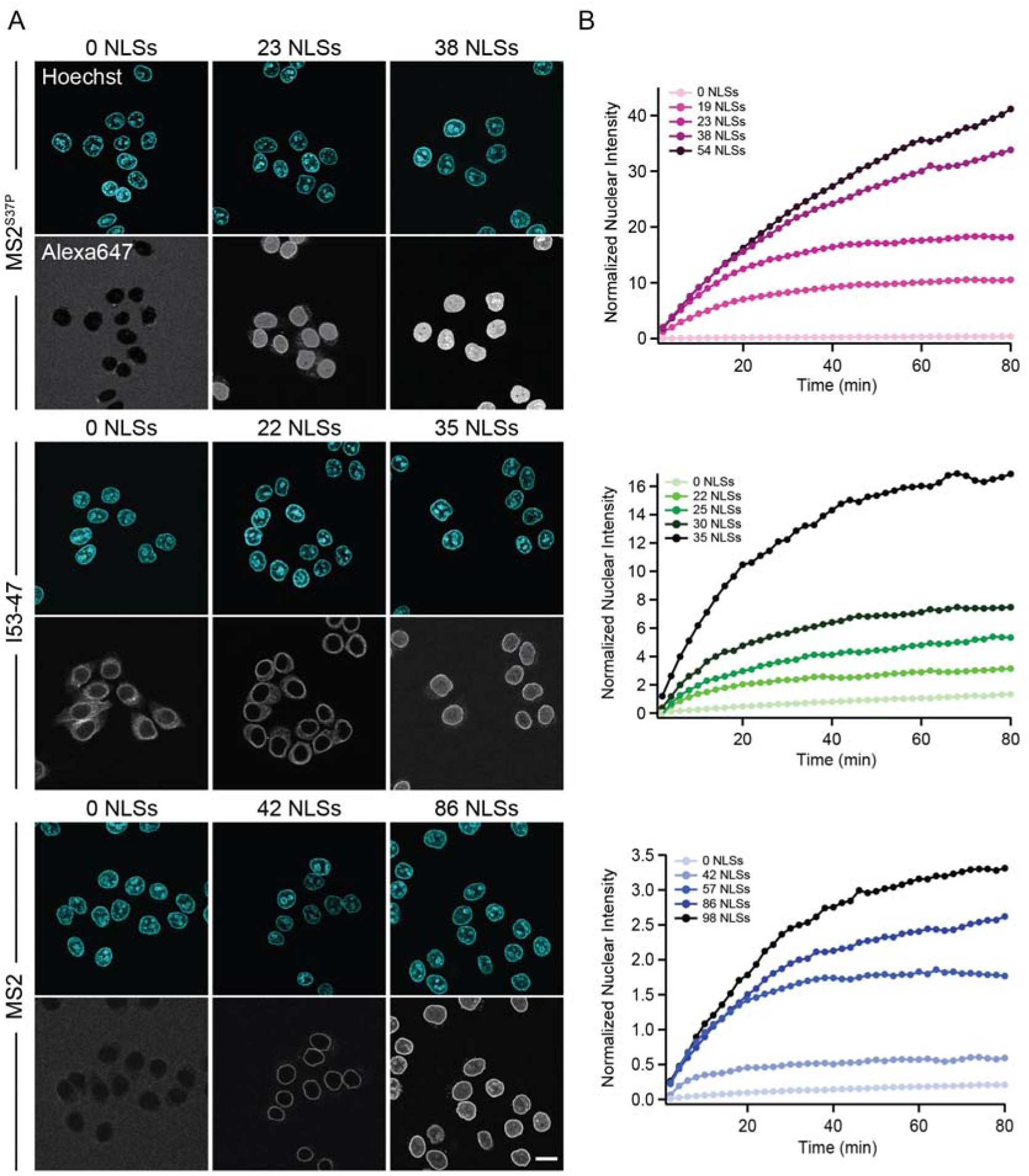
The import kinetic of large cargoes is tuned by the NLS number. (A) Confocal images of nuclear import of the different large cargoes. Cells were incubated for up to 1.5 h with capsids tagged with different number of NLS peptides on their surface. All cargoes displayed a distinct NLS-dependent behaviour. The scale bar corresponds to 20 μm. (B) Representative nuclear import traces for the three large cargoes labelled with increasing amount of NLS peptides. The nuclear intensities were normalized according to capsid brightness estimated from FCS and are plotted after subtracting the initial offset A determined by the fit.

### Modified HBV capsids are targeted to NPCs but do not accumulate into the nucleoplasm

We next subjected HBV capsids through the same pipeline. The maximum #NLSs labelling per capsid we achieved was 120, thus 50% of all capsid monomers got labelled. The capsids were targeted to the nuclear envelope, however no bulk nuclear import could be detected (**Figure 4**, first row). As we were not able to further increase the #NLS with our chemical labelling strategies and we wondered whether 120 NLSs might still be insufficient, we resorted to genetic tools to introduce one NLS per capsid monomer, thus 240 NLSs in total on the capsid. To do this, we designed a capsid based on the SplitCore construct (Walker et al., 2011), in which a core-GFP fusion protein is split into two halves that self-assemble before forming the capsid. This exposes a free C terminus, which we exploited to introduce an NLS. Also for this capsid, we did not observe any bulk import. However, the slightly increased size due to the GFP could potentially push this capsid over the maximum NPC transport limit. We thus tested another strategy, and introduced an NLS into an exposed capsid loop (**Figure 4**, last row). Again, no functional bulk import could be observed. EM showed that the engineered capsids are less homogenous, but still a large number of intact capsid was observed. Hence we conclude that none of the tested HBV capsids constructs can functionally be enriched in the nucleus. As the chances that our different careful modifications rendered the HBV capsid transport-incompetent seem rather low, our data does not agree well with a view that HBV capsids can pass through the NPC and enter the nucleus intact. Rather it is in line with studies that suggest that only the mature infectious virus can enter the NPC (Kann et al., 1999; Rabe et al., 2003), as those can be disassembled at the NPC to enter the nucleus. This is consistent with EM data on intact HBV capsids that show that they can enter the NPC barrier, and thus our data does not exclude this finding (Panté and Kann, 2002) (indeed, we do see strong NE accumulation). Note, that in contrast to the inside of the NPC, negative stain EM of permeabilised cells yields an ambiguous staining pattern in the nucleus, in which chromatin and capsid structures are not easily distinguishable. Collectively this suggests that 36 nm capsids might be able to enter the NPC barrier, but are too large to pass the NPC intact into the nucleus (i.e. undock or release). We thus focus in the following on our global quantitative analysis of the three capsids, for which satisfying conditions of functional import could be experimentally identified.

**Figure 4:**
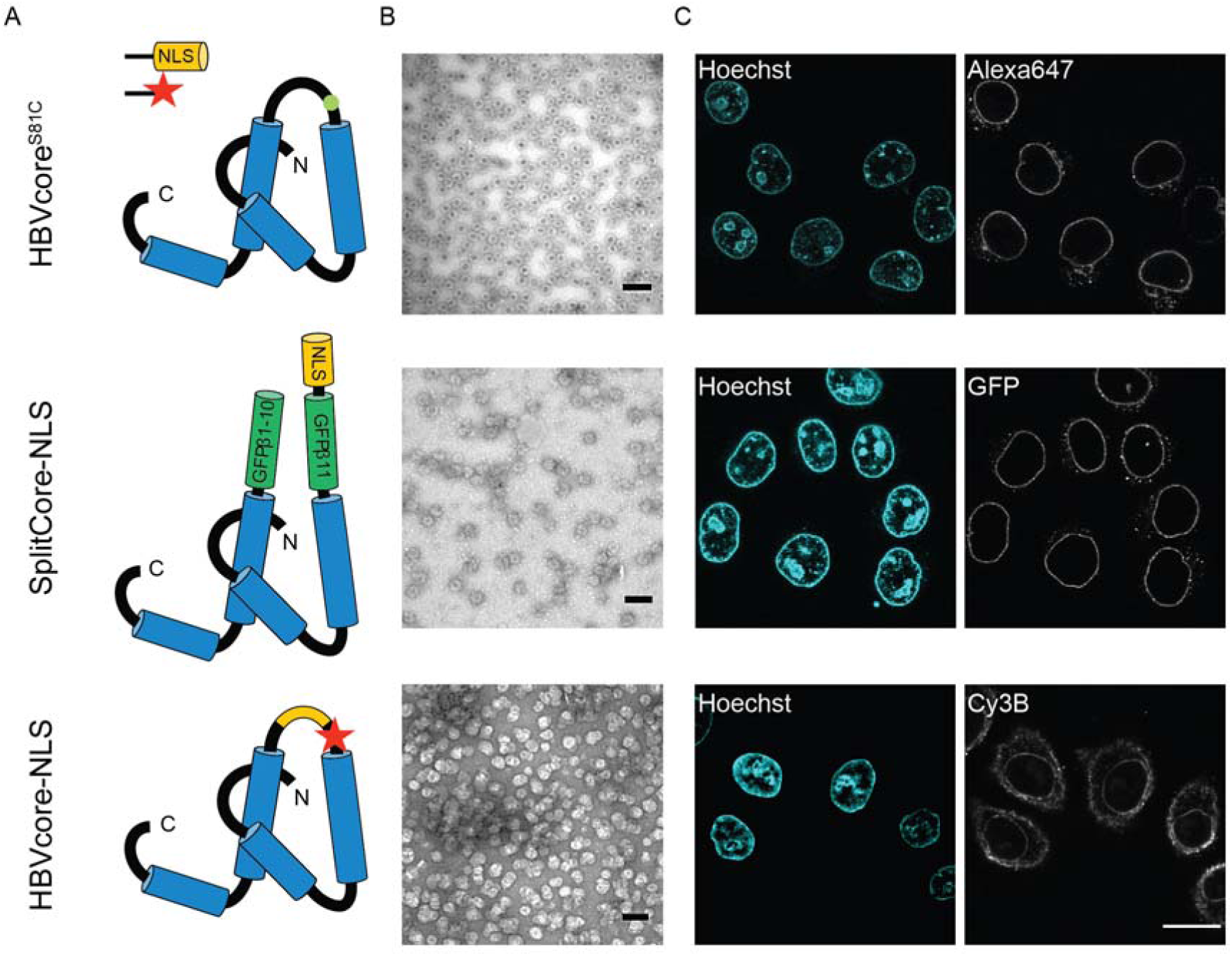
The Hepatitis B capsid is not imported in the nucleus of permeabilised cells. Following the same labelling approach as described in Figure 1, HBV capsids with up to 120 NLSs were generated (first row). In order to test capsids with a higher number of NLSs exposed on the surface, we designed two additional versions of the HBV core protein with a direct NLS insertion (cartoons in second and third row). All capsids were targeted to the nuclear envelope but did not give rise to bulk nuclear accumulation in import experiments using permeabilised cells. (A) Schematic representations of the different HBV core protein constructs. (B) EM images of the purified capsids. The scale bar corresponds to 100 nm. (C) Confocal images of capsid import experiments after 1.5 h. The scale bar corresponds to 20 μm.

### Global quantitative analysis of nuclear import in relation to cargo size and #NLSs

All nuclear import curves were fitted with a monoexponential kinetic model, with parameters I_MAX_ and T_1/2_. Overall, the two parameters correlated with each other (Supplementary Figure S3 and Table S1 for details), and thus we focus subsequent analysis on I_MAX_ which informs on the total number of imported capsids. In **Figure 5A** (left column) we plot the dependence of I_MAX_ on the #NLSs for all capsids. Cargoes with no NLS are plotted at 0 #NLSs but were excluded from the fit. For all capsids, we detected a linear increase of nuclear import with the #NLSs mounted onto the cargo. The ability to tune the #NLSs enables also to identify cut off limits and conditions in which bulk import can clearly happen. An extrapolation to I_MAX_ =0 yields an effective cut off limit of minimum #NLSs for import to occur. This was ≈10 NLSs for both the MS2^S37P^ and I53-47 capsids and ≈30 NLSs for MS2. The slope also varied dramatically depending on capsid size, with a slope of 1.2 for the 17 nm capsid (MS2^S37P^), 0.4 for the 23 nm (I53-47) and 0.04 for the 27 nm one (MS2).

**Figure 5:**
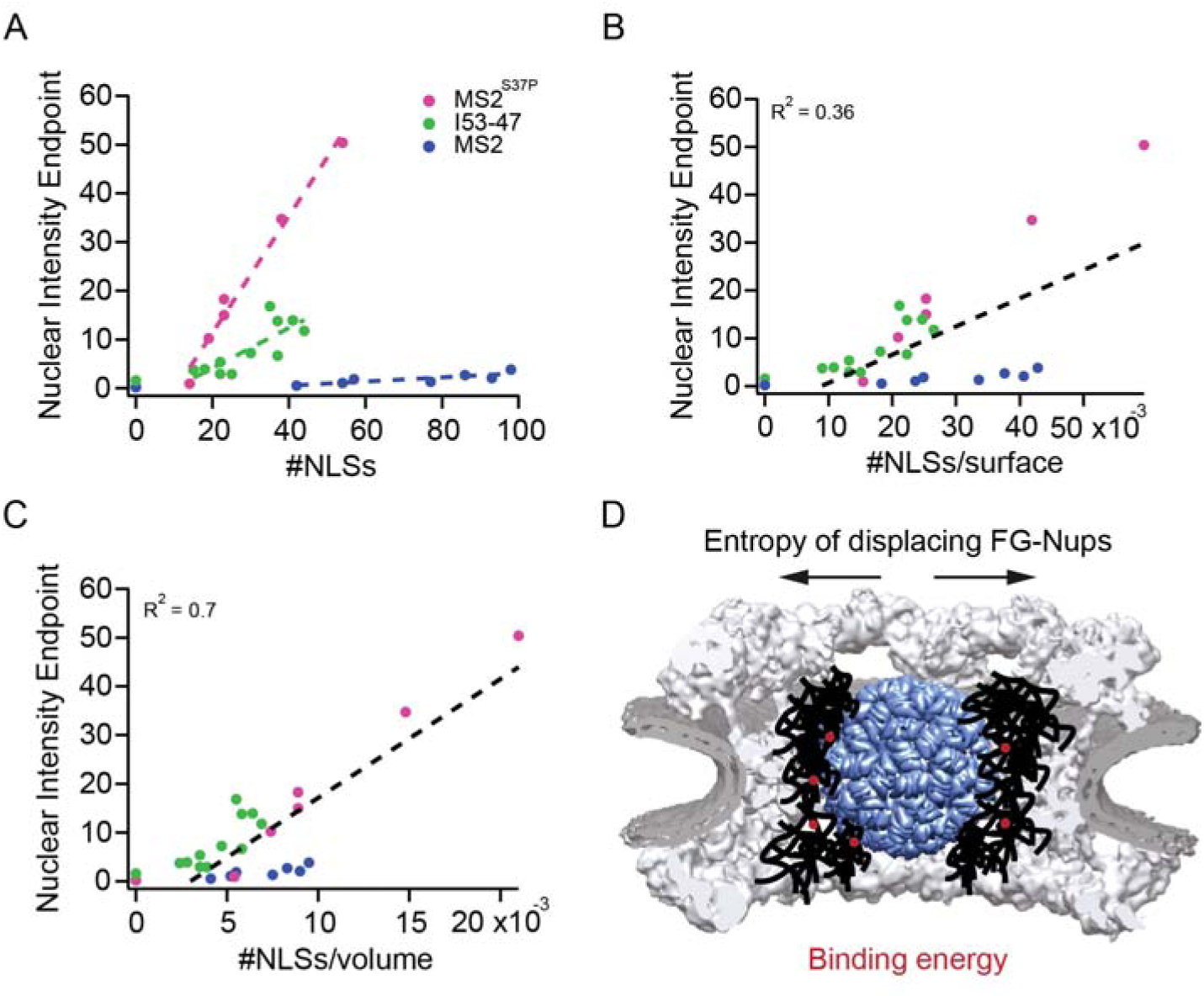
Effect of cargo size and number of NLSs on import efficiency and kinetics. The nuclear import kinetic curves were fitted with an inverse exponential function *I(t) = A* + *I*_*MAX*_ (1 – *e*^−*kt*^). (A) The plot on the left displays the nuclear intensity endpoint I_MAX_ (readout of import efficiency) for the different cargoes, in relation to the #NLSs coupled to the capsid. (B,C) correspond to the rescaled data, where we divide the #NLSs by the cargo surface and volume, respectively. (D) Cartoon of the determinants for large cargo import: the entropic cost of inserting a large spherical object in the dense FG-Nup barrier has to be compensated by binding via multiple NTRs (binding sites represented in red, NTRs omitted for simplicity). NPC scaffold structure from ref. (von Appen et al., 2015).

Tu et al previously reported a single molecule study of β-galactosidase which can have up to 4 NLS. This cylinder shaped molecule is 18 nm at its longest axis, similar to the MS2^S37P^. However, its smaller axis is 9 nm, which can explain well why for this substrate 4 NLS might be enough for import (Tu et al., 2013). The change in I _MAX_ and cut off value between MS2^S37P^ and the 6 nm larger (and structurally completely different) I53-47 appears much less dramatic as the difference between those two and the only 4 nm larger MS2 capsid. As the MS2 and small MS2^S37P^ share surface properties, since the individual core unit is nearly the same, it is suggested that surface properties of the capsids are not the major determinants for import (see Supplementary **Figure S4** for a comparative image of capsid surface properties). In fact, the substantial cargo decoration with Importins results in at least 1/3 of the capsid surface being shielded in conditions of minimum import cut off (Supplementary Figure S4). This probably distinguishes such large cargoes with multiple NLSs from much smaller cargoes with one or two NLSs, where surface differences can play a more substantial role (Frey et al., 2018).

The requirement of needing more than 10 NLSs to import MS2^S37P^ and I53-47 and more than 30 NLSs for MS2 provides important information on the energetic cost needed to immerse a large “spherical object” into the highly crowded NPC permeability barrier. In times where the number of computational models for NPC transport is increasing, but not necessarily converging, we hope these experimental values might help to benchmark different hypotheses.

We also attempted to identify common physical parameters of large cargo import without the need for a computational simulation. There are two simple and sound options on what might be a key parameter for an Importin-covered sphere to enter an FG-Nup barrier. In **Figure 5** we rescale our data once by surface and once by volume, leading to a plot of I_MAX_ vs #NLS/surface of the capsid and I_MAX_ vs #NLS/volume of the capsid respectively. While the linear fit is not perfect for the volume rescaling (R^2^ ≈0.7) it is better than for the square dependence (R^2^ ≈0.4). We thus cannot claim that a pure rescaling by volume captures all aspects of the data, and it would be informative to have more experiments, ideally from many more large structures. However, not many large cargoes appear suitable to this type of study, we are left with the total of 30 distinct samples we studied in this work. Those together point again to the fact that for large cargo transport, cargo volume to NLS ratio can capture essential aspects of large cargo shuttling through the NPC. To illustrate this process, we make a simple toy model calculation (**Figure 5D** for schematic representation) assuming that NTR binding contributes to reducing the energy barrier imposed by the entropic cost needed to displace FG-Nups. Based on the rough number of 10 FG-binding spots on the NTR surface (Isgro and Schulten, 2005), each Importinβ contributes roughly 10kT (Tu et al., 2013). For a bulk import cut off of 30 NLSs this translates into a total binding energy of 300kT per MS2 capsid. This is needed to move around ≈ 1 MDa of mass which can be extrapolated from the known capsid volume of ≈10000 nm^3^ and average FG-Nup concentration in the NPC of ≈2 mM (Frey and Gorlich, 2007). Note, on how in the cartoon in **Figure 5** a single MS2 capsid already occupies 1/3 of the estimated volume of the central NPC channel (Isgro and Schulten, 2005). The energetic cost required might be further lowered by additional mechanisms that enable the entire NPC scaffold to dilate, a hypothesis supported also by multiple evidences for tentative hinge elements in the NPC scaffold structures (Bui et al., 2013; Kelley et al., 2015). Future studies using our cargo substrates and time resolved high resolution measurements could provide further insights into the individual kinetic steps of NPC binding, barrier passage and undocking and how those link to FG-Nup and potentially scaffold dynamics in the NPC.

## Materials and methods

### Large cargo expression and purification

#### MS2 and MS2^S37P^ capsids

A colony of *E. coli* BL21 (DE3) AI cells containing the pBAD_MS2_Coat Protein-(1-393) or the pBAD_MS2_Coat Protein-(1-393)_S37P plasmids was inoculated in LB medium containing 50 μg/mL ampicillin. The culture was grown overnight shaking at 37°C (180 rpm) and then used at a 1:100 dilution to inoculate an expression culture in LB medium. Protein expression was induced at OD_600_=0.6-0.7 by adding 0.02% arabinose and carried out at 37°C shaking (180 rpm), for 4 hours. Cells were harvested by centrifugation at 4500 rpm, for 20 min, at 4°C. For purification, pellets were resuspended in an equal volume of lysis buffer (10 mM Tris pH 7.5, 100 mM NaCl, 5 mM DTT, 1 mM MgCl_2_, 1 mM PMSF) and lysed through 3-4 rounds in a microfluidizer, at 4 °C. The lysate was incubated with 0.2% PEI (Polyethylenimine) for 1 hour, on ice and then clarified by centrifugation at 10000 rpm, for 30 min. A saturated solution of (NH_4_)_2_SO_4_ was added at 4°C drop-wise to the clear lysate under continuous mild stirring up to 25% of ammonium sulphate. After 1 hour, the lysate was spun down in a centrifuge at 10000 rpm, for 30 min. The supernatant was discarded, and the pellets were gently resuspended with 10-20 mL of lysis buffer on a rotator, at room temperature. The lysate was then centrifuged at 10000 rpm, for 30 min and the clear supernatant was collected. The supernatant was cleared using the KrosFlo system (SpectrumLabs) with a 0.2 μm cutoff membrane to remove large impurities. The membrane permeate containing the cleared sample was collected on ice. In order to maximize protein recovery, the remaining supernatant was washed with 50 mL of lysis buffer and the permeate was pooled with the previously collected one. The sample was then concentrated using the KrosFlo with a 500 kDa cutoff membrane (for the smaller MS2^S37P^ capsid, a 30 kDa cutoff was used).

#### I53-47 capsids

A colony of *E. coli* BL21 (DE3) AI cells containing the pET29b(+)_I53-47A.1-B.3_D43C plasmid was inoculated in LB medium containing 50 μg/mL kanamycin. The culture was grown overnight shaking at 37°C (180rpm) and then used at a 1:100 dilution to inoculate an expression culture in LB medium. Protein expression was induced at OD_600_=0.8 by adding 1 mM IPTG and carried out at 37°C shaking (180 rpm), for 3 hours. Cells were harvested by centrifugation at 4500 rpm, for 20 min, at 4°C. The purification procedure was adapted from (Bale et al., 2016). Pellets were resuspended in two pellet volumes of lysis buffer (25 mM Tris pH 8.0, 250 mM NaCl, 20 mM imidazole, 1 mM PMSF, 0.2 mM TCEP), sieved to remove clumps and supplemented with 1 mg/mL lysozyme and DNAse. Cells were lysed by sonication on ice, and the lysate was clarified by centrifugation at 24000 g, for 35 min, at 4°C. The clear lysate was incubated with Ni-beads (1 mL/L expression) for 1-2 hours, at 4°C under gentle rotation. Ni-beads with lysate were poured in a polypropylene (PP) column and the flow through (FT) was collected. Ni-beads were washed three times with 20 mL of lysis buffer followed by elution with 5 mL of elution buffer, containing 500 mM imidazole. The elution was immediately supplemented with 5 mM EDTA to prevent Ni-mediated aggregation of the sample. The buffer of the protein was then exchanged to dialysis buffer (25 mM Tris pH 8.0, 150 mM NaCl, 0.2 mM TCEP), at 4°C. After dialysis, the protein was transferred to a new tube and spun down for 10 min, at 5000 rpm, at 4°C, in order to remove any precipitation. The protein was concentrated using the KrosFlo with a 100 kDa cutoff membrane, which also helps removing any remaining unassembled capsid proteins. After concentrating down to 3-4 mL of volume, the sample was washed with 50 mL of fresh dialysis buffer using the continuous buffer exchange mode of the KrosFlo.

#### HBV capsids

A colony of *E. coli* BL21 (DE3) AI cells containing the desired HBV plasmid was inoculated in TB medium containing 50 μg/mL ampicillin. The culture was grown overnight shaking at 37 °C (180rpm) and then used at a 1:100 dilution to inoculate an expression culture in LB medium. Protein expression was induced at OD_600_=0.8-1 by adding 0.02% arabinose and carried out at 20 °C shaking (180 rpm) overnight. Cells were harvested by centrifugation at 4500 rpm, for 20 min, at 4°C. The purification procedure was adapted from (Walker et al., 2011). Pellets were resuspended in one pellet volume of lysis buffer (25 mM Tris pH 7.5, 500 mM NaCl, 0.2 mM TCEP, 10 mM CHAPS) and lysed by sonication 3×30 seconds, on ice. The lysate was spun down at 10000 rpm, for 10 min. The cleared supernatant was then loaded on a step gradient 10-60% sucrose obtained by mixing lysis and sucrose buffers (25 mM Tris pH 7.5, 500 mM NaCl, 0.2 mM TCEP, 10 mM CHAPS, 60% sucrose) in appropriate ratios and by carefully layering the different percentage buffers into ultracentrifugation tubes. The lysate was then subjected to ultra-centrifugation at 28000 rpm, for 3.5 hours at 4°C. Fractions of 2 mL were collected by gravity, by puncturing the ultracentrifugation tube from the bottom. Fractions containing the capsids were pooled and concentrated using the KrosFlo with a 500 kDa cutoff membrane.

### Large cargo maleimide labelling and characterization

Purified capsids were labelled via maleimide chemistry to couple a fluorescent dye and NLS peptide to the exposed cysteines. The dye (AlexaFluor647 maleimide, Invitrogen) and NLS peptide (Maleimide-GGGGKTGRLESTPPKKKRKVEDSA, PSL Peptide Specialty Laboratories) were stored at −80°C and freshly resuspended in anhydrous DMSO. The capsids were incubated with different molar excesses of dye and NLS peptide according to the desired degree of labelling for 1-2 hours, at room temperature. The reaction was then quenched by adding 10 mM DTT and the protein was spun down at 10000 rpm, for 10 min, at 4°C to remove any precipitation. The excess dye was removed by loading the capsid sample on a HiPrep Sephacryl 16/60 size exclusion column (GE Healthcare), using the appropriate buffer (for MS2 and MS2^S37P^: 10 mM Tris pH 7.5, 100 mM NaCl, 5 mM DTT; for I53-47: 25 mM Tris pH 8.0, 150 mM NaCl, 1 mM DTT and for HBV: 25 mM Tris pH 7.5, 500 mM NaCl, 0.2 mM TCEP, 10 mM CHAPS, 10% sucrose). Relevant fractions containing the labelled capsids were then pooled and concentrated using the KrosFlo. For long-term storage at −80°C, the sample was supplemented with 25% glycerol (30% sucrose for HBV) and either flash-frozen with liquid nitrogen or directly transferred to the −80°C freezer (for I53-47 capsids). The ratio of capsid monomers tagged with NLS peptide was quantified by the gel band ratio on a SDS PAGE gel with Coomassie staining, as the labelled monomers migrate differently due to their increased molecular weight.

### Electron microscopy

Capsid integrity was confirmed by imaging the samples with an electron microscope using negative staining. Carbon-coated 300 meshes Quantifoil Cu grids were glow-discharged for 10 s in a vacuum chamber. Then, a 3 μL drop of sample was adsorbed on a grid for 2 min, blotted with Whatman’s filter paper and washed 3 times with sample buffer, then 3 times with a solution of 2% uranyl acetate. Once the grids were dry, the sample was imaged using a Morgagni 268 microscope (FEI).

### Dynamic light scattering

Dynamic light scattering (DLS) measurements to quantify the hydrodynamic radius of capsids and test for sample aggregation or disassembly were performed on a Zetasizer Nano (Malvern). Samples were diluted to a final concentration of 0.5 μM in filtered TB and spun down for 10 min at 10000 rpm prior to each measurement. For each sample, at least 10 measurements were acquired, using a 12 μl quartz cuvette. Count rates per second were typically higher than 200 kcps, and the polydispersity index was below 0.2, indicating a monodisperse solution. Data were analysed using the Malvern software, using the Multiple Narrow Bands fitting algorithm and Refractive Index and Absorption settings for proteins.

### Fluorescence correlation spectroscopy

Fluorescence correlation spectroscopy (FCS) was used to characterize the large cargoes and quantify their concentration and brightness (#dyes/capsid). FCS experiments were carried out on a custom-built multiparameter spectrometer confocal setup, equipped with a 60x water objective (NA = 1.27). The capsid samples were diluted in freshly filtered 1XTB and spun down for 10 min at 10.000 rpm at 4°C prior to the start of the experiment. FCS measurements were carried out in 8-well Lab-Tek, which had been pre-incubated for 30 minutes with a solution of 1 mg/ml BSA to prevent sample sticking. For each sample, at least 10 FCS curves of 30 seconds each were acquired. Low power (1-5 μW) was used to avoid bleaching of the samples during their diffusion through the confocal volume. A calibration FCS measurement with a free dye solution was carried out every 2-3 samples to measure the structural parameter and confirm the stability of the setup. Data analysis was performed with SymphoTime software. Autocorrelation curves were computed for lag times between 0.0001 and 1000 ms and fitted with a diffusion model. Capsid brightness was calculated by dividing the measured particle brightness by the measured brightness of a calibration dye solution at the same laser power settings.

### Nuclear import assays

HeLa Kyoto cells were cultured at 37°C, 5% CO_2_ atmosphere in Dulbecco’s modified Eagle’s medium with 1 g/mL glucose (Gibco 31885023) supplemented with 1% penicillin-streptomycin (Sigma P0781), 1% L-Glutamine (Sigma G7513) and 10% FBS (Sigma F7524). The cells were passaged every 2-3 days up to maximum of 15-17 passages. Cells were seeded 1 or 2 days prior the experiment at low density (10000-12000 cells per well) in a glass bottom 8-well Lab-Tek II chambered coverglass (Thermo Scientific Nunc, 155383).

Cells for transport assays were stained with 100 nM MitoTracker green (Invitrogen, M7514) in growth medium for 30 min, at 37°C, 5% CO_2_. For nuclear staining, cells were rinsed once with PBS and incubated for 10 min, at room temperature with 20 nM Hoechst 33342 (Sigma, B2261). Cells were then washed once with transport buffer (1XTB: 20 mM Hepes, 110 mM KOAc, 5 mM NaOAc, 2 mM MgOAc, 1 mM EGTA, pH 7.3 adjusted with KOH) and permeabilised by incubation for 10 min, at room temperature with digitonin (40 μg/mL). Cells were then washed 3 times with 1XTB supplied with 5 mg/mL PEG 6000 to avoid osmotic shock. After the final wash, excess buffer was removed and the transport mix was quickly added to the cells to start the experiment. The final transport mix was composed of 1 μM Importinα, 1 μM Importinß, 4 μM RanGDP, 2 μM NTF2, 2 mM GTP and 8 nM capsid cargo. In order to allow the import complex to form, the cargo was first pre-incubated with Importinß and Importinα on ice for at least 10 minutes, then the rest of the transport mix was added and the solution was spun down for 10 min at 10000 rpm to remove any aggregates. Each experiment was performed side-by-side with control cells incubated with fluorescently labelled 70 kDa Dextran (Sigma 53471) to confirm nuclear envelope intactness throughout the whole experiment.

### Confocal fluorescence microscopy

Time-lapse confocal imaging of nuclear import was performed on an Olympus FLUOVIEW FV3000 scanning confocal microscope, using a 40x air objective (NA = 0.95). An automated multi-position acquisition was carried out, where 12 different regions (typically containing 10 cells each) were imaged in two different wells. Three channels were recorded at each time step, using the 405 nm (Hoechst), 488 nm (Mitotracker) and 640 nm (cargo) laser lines for excitation. Images were acquired every 2 minutes for 80-90 minutes, using continuous autofocusing with Z-drift compensation to ensure imaging stability.

### Image and data analysis

Results of the time-lapse import experiments were analysed with a custom-written Fiji script. The Hoechst and Mitotracker channels were used to generate reference masks for the nucleus, nuclear envelope and cytoplasm at each time point. Briefly, the two images were pre-processed with Gaussian blur to aid in area segmentation, and then thresholded. The nuclear mask was eroded three times to remove contributions coming from the nuclear envelope, and the envelope mask was generated by subtracting the eroded mask from the non-eroded one. The final masks were then used to extract the average intensity of cargo signal in the different areas of interest. Final data analysis and plotting was performed in IgorPro (Wavemetrics). Fluorescence intensities were background-corrected, rescaled according to the capsid brightness and fitted to an inverse exponential function I(t)=A+I_max_(1-e^-kt^). The time to half saturation was calculated as T_1/2_ =ln2/I_max_.

## Online supplemental materials

Figure S1. I53-50 capsid.

Figure S2. Control experiments in permeabilised cells. Figure S3. Entire import kinetic dataset.

Table S1. Parameters from import kinetics fits.

Figure S4. Comparison of large cargo surface properties.

## Acknowledgments

We are particularly grateful to Joana Caria for her passionate help in preparing this work. We also thank Gemma Estrada Girona and Miao Yu for their contributions to this work. We thank SFB 1129 for generous funding (Projektnummer 240245660 funded by DFG, German Research Foundation). We thank the Baker lab for providing plasmids for their artificial capsid structure and Michael Nassal for HBV constructs. We thank Anton Zilman and Ulrich Schwarz for insightful discussion. We also thank the ALMF and EM facilities at EMBL and the mechanical workshop.

