## Supplementary Material for "Molecular determinants of large cargo transport into the nucleus"

**Supplemental materials**

Figure S1. **I53-50 capsid.** Analog to main text Figure 1 (A) shows I53-50 capsid structures rendered in Chimera (Pettersen et al., 2004) (left) and EM image of the purified sample (right). The scale bar corresponds to 50 nm. (B) SDS-PAGE gel of I53-50 capsids labelled with Alexa647. The different columns correspond to fractions from the main peak of the size exclusion column. Note, how fluorescent labelling is observed in both chain A (top band) and chain B (bottom band), while the labelling aimed only to label an inserted reactive cysteine residue in chain B. This labelling ambiguity could result in an inaccurate estimate of the #NLSs coupled to the capsid surface, therefore we decided to exclude this capsid from further analysis.

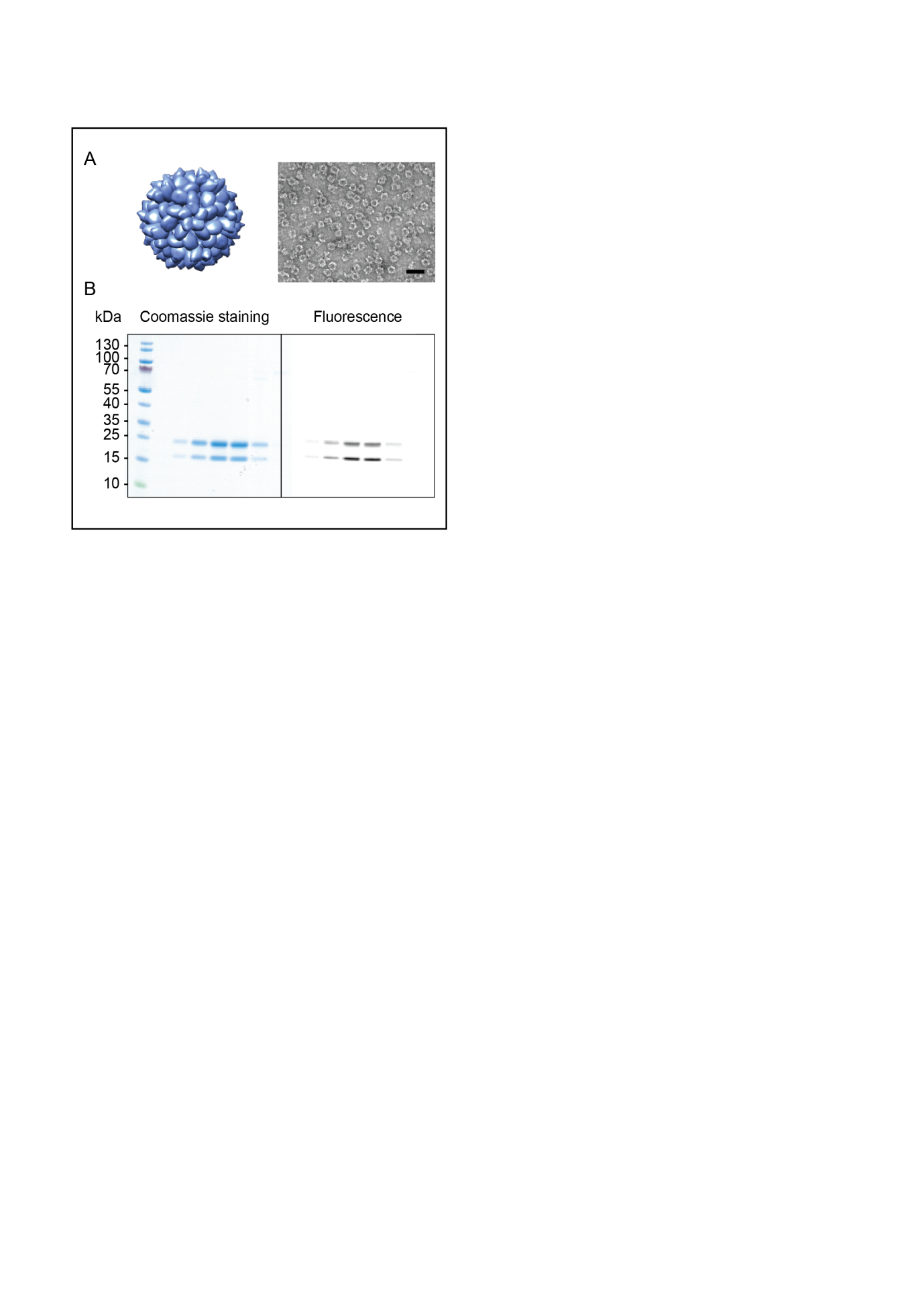

Figure S2**. Control experiments in permeabilised cells.** In order to validate the transport assays in permeabilised cells, we performed several control experiments to rule out the possibility that the measured kinetics were e.g. influenced by depletion of components in the transport mix during the course of the experiment. Panel A shows a comparison of the same MS2^S37P^ sample measured in normal conditions, with a 5-fold excess of Importinα, with a 2-fold excess of GTP and with addition of an energy regeneration system to the transport mix (0.1 mM ATP, 4 mM creatine phosphate and 20U/ml creatine kinase). For all cases, capsid import did not change substantially compared to the typical variability in these experiments (the shaded area represents the standard deviation of intensity over 12 different areas acquired). In order to further exclude issues with recycling of transport mix components over time, we purified CAS (the protein responsible for shuttling Importinα back into the cytoplasm (Sun et al., 2013)) and tested its effect on the transport of a MS2^S37P^ sample with 23 NLSs. As can be seen in panel B, including 1 µM CAS in the transport mix, only minor differences in the kinetics of the two experiments were observed.

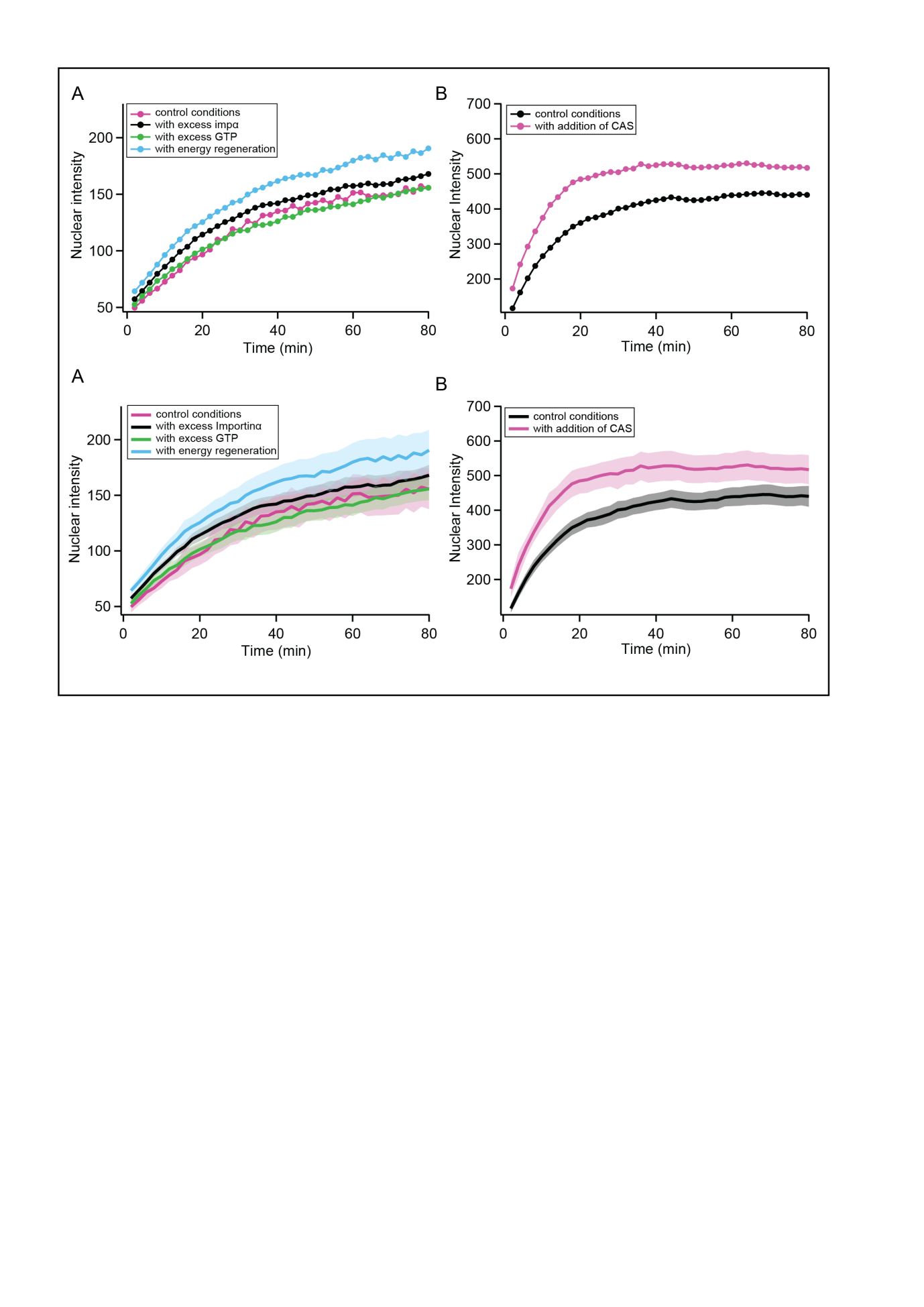

Figure S3. **Entire import kinetic dataset.** Corresponding to main Figure 3, here we show all measured kinetics for the MS2^S37P^ (A), I53-47 (B) and MS2 (C) capsids. The traces represent the average nuclear intensity measured in 12 different areas, background-subtracted and corrected to account for the different sample brightness. Overlaid on top of the traces, we show the inverse exponential fits used to extract the I_MAX_. As sample adhesion to the coverslip/unspecific sticking increased for some samples over the course of the experiment, giving rise to a linear background increase, we limited the fitting to the first 40 minutes to extract more accurate kinetics. The most reliable parameter to extract from the fit was I_max_, and thus we chose to focus on the analysis of this one further. This parameter correlates to a large extent with the T_1/2_ (R^2^ ≈ 0.9 for MS2^S37P^ and MS2 as well as 0.4 for I53-47).

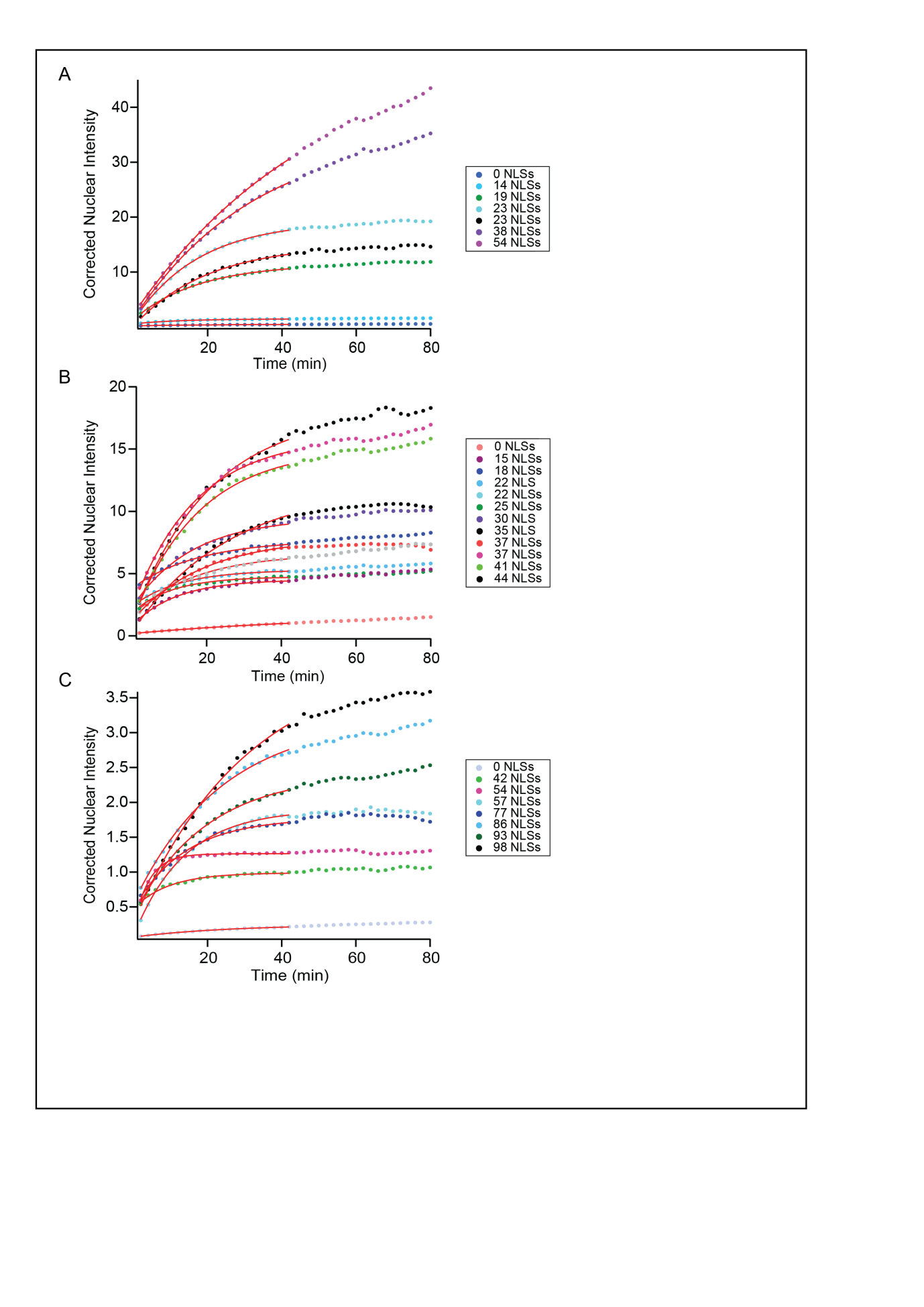

Table S1. **Parameters from import kinetics fits.** Here we list all parameters extracted from fitting the import traces with an inverse exponential I(t) = A + I_MAX_ (1-e^-kt^). T_1/2_ is calculated as ln2/k.

|  | **sample** | **#NLSs** | **A** | **I_MAX_** | **k** | **T_1/2_** |
| --- | --- | --- | --- | --- | --- | --- |
| MS2^S37P^ | 1 | 0 | 0.21 | 0.28 | 0.050 | 13.86 |
|  | 2 | 14 | 0.57 | 0.91 | 0.078 | 8.89 |
|  | 3 | 19 | 1.34 | 10.18 | 0.057 | 12.16 |
|  | 4 | 23 | 1.08 | 18.24 | 0.057 | 12.25 |
|  | 5 | 23 | 0.11 | 14.97 | 0.050 | 13.86 |
|  | 6 | 38 | 1.41 | 34.69 | 0.030 | 23.10 |
|  | 7 | 54 | 2.31 | 50.39 | 0.020 | 35.36 |
| I53-47 | 1 | 0 | 0.18 | 1.58 | 0.018 | 38.51 |
|  | 2 | 15 | 0.80 | 3.71 | 0.085 | 8.15 |
|  | 3 | 18 | 3.81 | 3.89 | 0.056 | 12.36 |
|  | 4 | 22 | 2.34 | 2.95 | 0.080 | 8.71 |
|  | 5 | 22 | 2.01 | 5.35 | 0.060 | 11.61 |
|  | 6 | 25 | 1.83 | 2.91 | 0.094 | 7.37 |
|  | 7 | 30 | 2.27 | 7.22 | 0.063 | 11.02 |
|  | 8 | 35 | 1.18 | 16.82 | 0.048 | 14.44 |
|  | 9 | 37 | 1.21 | 6.67 | 0.053 | 13.15 |
|  | 10 | 37 | 2.16 | 13.80 | 0.059 | 11.83 |
|  | 11 | 41 | 1.13 | 13.92 | 0.057 | 12.27 |
|  | 12 | 44 | 0.31 | 11.78 | 0.038 | 18.29 |
| MS2 | 1 | 0 | 0.07 | 0.18 | 0.038 | 18.48 |
|  | 2 | 42 | 0.47 | 0.52 | 0.106 | 6.54 |
|  | 3 | 54 | 0.19 | 1.07 | 0.241 | 2.88 |
|  | 4 | 57 | 0.07 | 1.83 | 0.074 | 9.43 |
|  | 5 | 77 | 0.47 | 1.31 | 0.072 | 9.67 |
|  | 6 | 86 | 0.55 | 2.67 | 0.042 | 16.54 |
|  | 7 | 93 | 0.38 | 2.04 | 0.051 | 13.56 |
|  | 8 | 98 | 0.27 | 3.82 | 0.033 | 21.26 |

Figure S4. **Comparison of large cargo surface properties.** (A) Coulombic surface coloring of the three kinetically investigated capsids, generated in Chimera (Pettersen et al., 2004). The color scale is in units of *kcal/(mol*e)*, where *e* is the charge of a single electron. (B) Hydrophobicity surface coloring generated in Chimera, the color scale refers to units in the Kyte-Doolittle hydrophobicity scale (Kyte and Doolittle, 1982). Despite the fact that capsids show a complex landscape of surface charge and hydrophobicity properties, these do not differ substantially between the three capsids. Additionally we consider that each Importinβ has a footprint of roughly 50 nm^2^. For the minimum #NLS needed for efficient bulk nuclear import of the different viruses (maintext Figure 5), taking into account that NLS and Importin form a 1:1 stochiometric complex, the lower boundary for surface shielding by Importin is roughly 50% for MS2^S37P^, 30% for I53-47 and 60% for MS2.

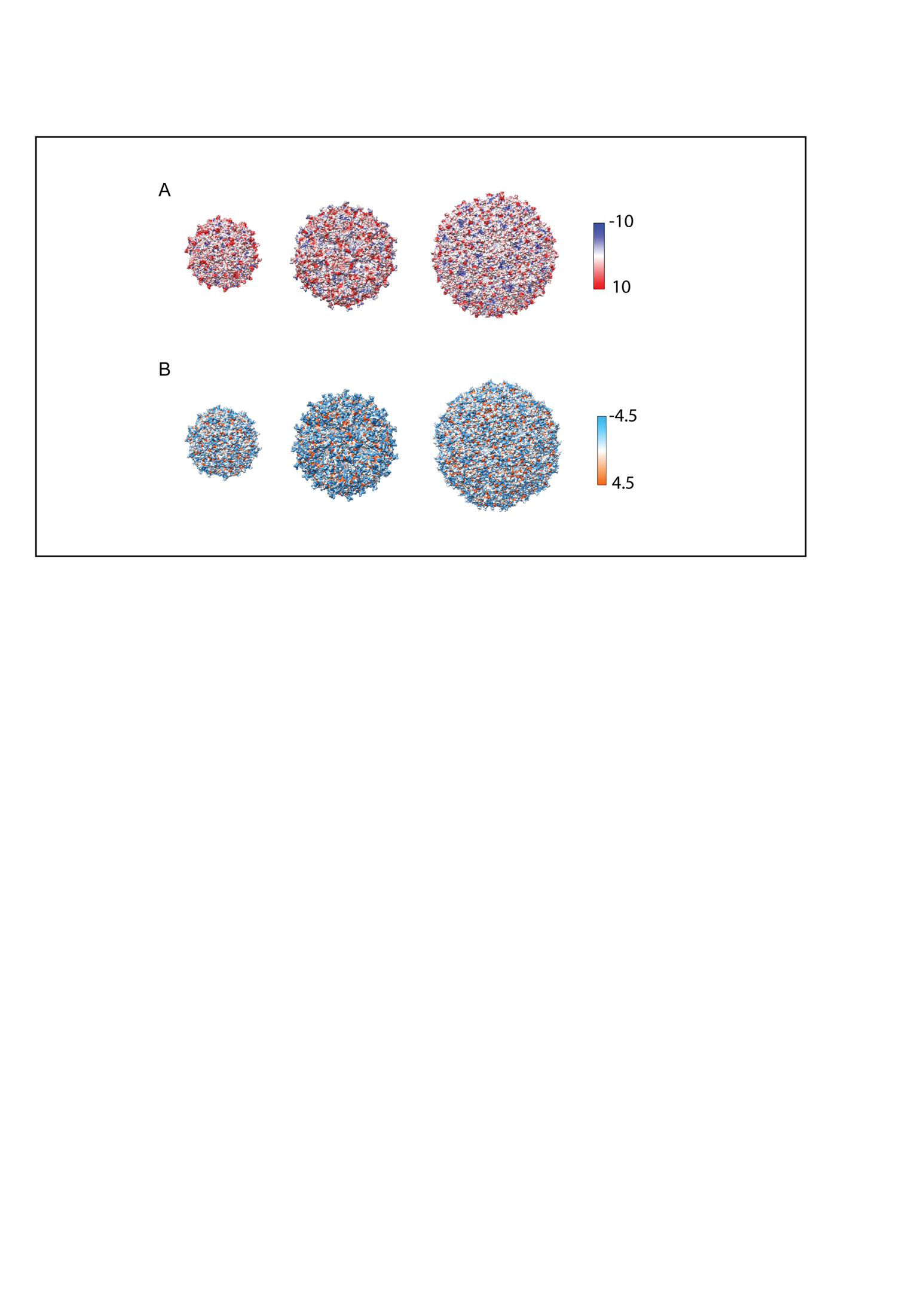
